# DrugLM: A Unified Framework to Enhance Drug-Target Interaction Predictions by Incorporating Textual Embeddings via Language Models

**DOI:** 10.1101/2025.07.09.657250

**Authors:** Tianyi Li, Zhengyu Fang, Xiaoge Zhang, Kaiyu Tang, Huiyuan Chen, Zhimeng Jiang, Tianxiang Zhao, Rong Xu, Feixiong Cheng, Xiao Li, Jing Li

## Abstract

**Motivation:** Accurate prediction of drug–target interactions (DTIs) is central to computational drug discovery, offering the potential to reduce experimental costs and accelerate development timelines. While existing deep learning approaches such as Graph Neural Networks and Transformers have shown promise, they often overlook the rich semantic information embedded in textual descriptions of drugs and targets. These descriptions encode critical biomedical knowledge, including mechanisms of action, biological pathways involved, and therapeutic effects of drugs, which can enhance DTI prediction performance.

**Results:** We introduce *DrugLM*, a unified framework that integrates embeddings derived from large language models (LLMs) into DTI-specific model architectures. DrugLM leverages textual descriptions of drugs and targets to generate semantic embeddings using a range of pretrained LLMs. These embeddings can be seamlessly incorporated into existing DTI models. We systematically evaluate multiple LLMs on benchmark DTI datasets and demonstrate strong performance even without fine-tuning. Moreover, supervised parameter-efficient fine-tuning of the LLMs further improves embedding quality, leading to enhanced prediction accuracy. Notably, a simple multilayer perceptron (MLP) using only LLM-derived embeddings surpasses several established DTI methods, underscoring the power of semantic features. Our findings highlight the practical value of integrating LLMs into DTI pipelines and offer a straightforward recipe for improved drug discovery: LLM embeddings of drugs and targets are both effective and easy to use.

**Availability:** Our code and dataset are available at https://github.com/ShPhoebus/DrugLM

## 1 Introduction

Traditional drug discovery is notoriously expensive, time-consuming, and prone to high failure rates (Mullard, 2014; Menden *et al*., 2019). Estimates suggest that the development of a single drug can cost between$314 million and$2.8 billion, with clinical phases spanning 8.2 to 10 years (Brown *et al*., 2021). To mitigate these challenges and reduce dependence on resource-intensive laboratory procedures, substantial efforts have been directed toward computational approaches, particularly those leveraging deep learning (Gawehn *et al*., 2015; Chen and Li, 2019b, Chen and Li, 2018). Among various computational drug discovery tasks, drug–target interaction (DTI) prediction plays a foundational role, offering mechanistic insights into pharmacological interventions (Wu *et al*., 2018; Chen *et al*., 2020a; Xiong *et al*., 2021).

Recent deep learning frameworks have achieved impressive results in DTI prediction (Jones *et al*., 2021; Chen and Li, 2020), especially those based on Graph Neural Networks (GNNs) (Yang *et al*., 2022; Xu *et al*., 2023, 2025; Fang *et al*., 2025) and Transformer-based models (Huang *et al*., 2020). For instance, GraphDTA (Nguyen *et al*., 2020) represents drug chemical structures as graphs and employs GNNs to estimate drug–target affinity. Similarly, MolTrans (Huang *et al*., 2020) adopts a Transformer-based architecture to capture interactions between molecular substructures, using linear representations of SMILES strings and amino acid sequences. While these models are effective, they primarily rely on data from a single modality: either one-dimensional string-based representations or two-dimensional graphs. They often overlook information from other modalities, such as three-dimensional structural information, or textual descriptions. More recently, there is a growing interest in integrating multimodal biological data, including three-dimensional structural information (Liu *et al*., 2025) and textual descriptions (Vazquez *et al*., 2011; Chen and Li, 2019a; Tarca *et al*., 2021), to further enhance DTI prediction performance.

Recently, language models (LMs) have emerged as powerful tools for applications in biology and chemistry (Guo *et al*., 2023; Lin *et al*., 2023; Hie *et al*., 2023; Zhao *et al*., 2025). For instance, BioT5 integrates molecular string representations with protein names, sequences, and structural data to train LMs on diverse prediction tasks (Pei *et al*., 2023). Similarly, DTI-LM employs LMs to encode protein amino acid sequences and drug SMILES strings for DTI prediction (Ahmed *et al*., 2024). However, existing approaches often overlook the rich semantic information embedded in unstructured textual descriptions of drugs and targets. These descriptions encapsulate critical biomedical knowledge, such as mechanisms of action, biological pathways involved, and therapeutic effects, which could potentially improve the accuracy of DTI prediction (Singhal *et al*., 2023).

In this work, we aim to incorporate prior knowledge of drugs and protein targets to facilitate DTI prediction. To this end, we propose *DrugLM*, a unified framework that integrates embeddings derived from large language models (LLMs) into DTI-specific architectures. DrugLM follows a two-stage design. In the first stage, it generates semantic embeddings of drugs and targets using their free-text descriptions and multiple pretrained LLMs. For this work, we select three LMs from the top performers on the MTEB leader board ^1^. In the second stage, these embeddings are seamlessly incorporated into downstream DTI prediction backbones. We include five widely used DTI prediction models (DeepConv-DTI (Lee *et al*., 2019), GraphDTA (Nguyen *et al*., 2020), NGCF (Wang *et al*., 2019), LightGCN (He *et al*., 2020), BACPI (Li *et al*., 2022)) and evaluate their performance with the three pretrained LMs on a curated drug–target dataset. Results show that they all perform better than the original models in terms of AUC and accuracy. Additionally, we apply parameter-efficient supervised fine-tuning on these LLM models, resulting in refined embeddings and notable improvements in DTI prediction accuracy. These results demonstrate the value of text-derived semantic features and the potential of LLMs to enhance DTI prediction pipelines.

In summary, our main contributions are as follows:

- **Incorporating textual semantics improves DTI prediction.** We demonstrate that integrating rich textual descriptions of drugs and targets significantly boosts the performance of various DTI models. Specifically, incorporating pretrained LM embeddings into the BACPI framework results in a 38.48% improvement over its baseline performance. Similarly, applying this approach to the GraphDTA framework yields a 12.21% increase in prediction accuracy.
- **Parameter-efficient fine-tuning enhances embedding quality** We show that supervised, parameter-efficient fine-tuning of language models leads to superior results compared to using pretrained models directly. This approach improves embedding quality via accelerated training. For example, within the GraphDTA framework, fine-tuning delivers an additional 8.96% performance gain over the non-fine-tuned counterpart.
- **DrugLM is a flexible and effective solution for DTI** Our proposed *DrugLM* framework can be seamlessly integrated into any existing DTI backbone. Notably, a simple MLP-based architecture using DrugLM embeddings outperforms several well-established DTI models. We offer a practical and effective recipe for DTI prediction: *LLM-derived embeddings of drugs and targets are surprisingly effective for drug discovery*.

## 2 Related Works

In this section, we review key computational approaches for drug–target interaction prediction and highlight recent advances in language models that have been applied to drug discovery tasks.

### 2.1 Computational Drug-Target Prediction

Numerous computational techniques have been developed to address the challenge of DTI prediction, aiming to reduce the high costs and time demands of experimental methods (Lee *et al*., 2019; Bai *et al*., 2023; Lim *et al*., 2019; Chen *et al*., 2020b,a; Chen and Li, 2019b; Nguyen *et al*., 2020; Yang *et al*., 2024; Wang *et al*., 2024, Wang *et al*., 2023b; Yan *et al*., 2023). Among these, deep learning approaches have gained significant attention for their ability to automatically extract informative features from raw inputs and capture complex non-linear relationships. For example, DeepConv-DTI (Lee *et al*., 2019) employs convolutional neural networks (CNNs) to learn local patterns from protein sequences and drug SMILES representations. GraphDTA (Nguyen *et al*., 2020) represents drug molecules as graphs and leverages graph neural networks (GNNs) to better capture topological and structural properties. These deep models have consistently outperformed traditional techniques such as matrix factorization (Chen *et al*., 2020a), owing to their capacity for hierarchical feature learning.

Graph-based collaborative filtering methods model drugs and targets as nodes in a bipartite interaction graph and learn their embeddings by propagating information across edges. Neural Graph Collaborative Filtering (NGCF) (Wang *et al*., 2019) injects both first- and higher-order collaborative signals into the embedding process via layer-wise message passing with nonlinear feature interaction, explicitly capturing complex connectivity patterns in the drug–target graph. LightGCN (He *et al*., 2020) simplifies this approach by removing feature transformations and activation functions, relying solely on normalized neighborhood aggregation and layer combination.

More recently, attention mechanisms and Transformer-based architectures have been introduced to further improve model expressiveness and interpretability. Bi-directional Attention-based Compound–Protein Interaction (BACPI) (Li *et al*., 2022) integrates a graph attention network to encode compound structures and a convolutional neural network to encode protein sequences, fusing them via a bi-directional attention module that lets atom- and residue-level features attend to each other. DrugBAN (Bai *et al*., 2023) incorporates a bilinear attention network to focus on meaningful drug substructures and protein residues, enhancing both prediction accuracy and biological interpretability. MolTrans (Huang *et al*., 2020) adopts a self-attention framework to model interactions between subcomponents of drugs and proteins, treating DTI prediction as a sequence-to-sequence translation task. TransformerCPI (Chen *et al*., 2020b) uses a Transformer encoder to capture contextual relationships within protein sequences and drug representations, achieving strong generalization to unseen interactions. Despite these advances, most existing models overlook the rich semantic information embedded in textual descriptions of drugs and targets. Such descriptions often encode valuable biomedical knowledge, including mechanisms of action, biological pathways involved, and therapeutic effects (Wang *et al*., 2023a; Zheng *et al*., 2024), which, if properly leveraged, could significantly enhance DTI prediction performance.

### 2.2 Language Models for Drug Discovery

LLMs have recently demonstrated remarkable capabilities in understanding scientific language and supporting a wide range of downstream tasks critical to drug discovery and development (Zheng *et al*., 2024; Nguyen and Grover, 2025; Wang *et al*., 2025). A prominent example is Geneformer (Theodoris *et al*., 2023), an LLM trained on 30 million single-cell transcriptomes. Geneformer has shown potential in disease modeling and was able to identify candidate therapeutic targets for cardiomyopathy through in silico gene deletion experiments.

In the context of DTI prediction, DTI-LM (Ahmed *et al*., 2024) employs ESM-2 (Lin *et al*., 2023) to encode protein amino acid sequences and ChemBERTa (Chithrananda *et al*., 2020) to represent drug SMILES strings, demonstrating how specialized protein and molecular LMs can be integrated for end-to-end drug–target interaction modeling.

In contrast to these approaches, our proposed **DrugLM** directly leverages large language models to derive embeddings from natural language descriptions of both drugs and targets. This enables a more flexible and semantically enriched representation, potentially enhancing the accuracy and interpretability of DTI predictions.

## 3 Materials and methods

### 3.1 Problem Setup

We formulate DTI prediction as a binary classification task, where the goal is to determine whether a given drug–target protein pair is likely to interact. Let the set of drugs be denotedSS by 𝒟 = *{d*^1^, *d*^2^, …, *d*^*m*^*}* and the set of target proteins by 𝒫 = *{p*^1^, *p*^2^, …, *p*^*n*^*}*, where *m* and *n* are the total numbers of drugs and targets, respectively. The drugs and targets can have additional information associated with them, for examples, drug chemical structures, target sequences and structures. With a slight abuse of notation, we assume that such information is encoded in 𝒟 and 𝒫. We further assume that each drug is associated with a natural language textual description, denoted as 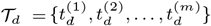. Similarly, each protein is also associated with a textual description, denoted as 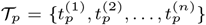.

These descriptions are free text in natural language and can encapsulate rich biomedical knowledge, including mechanisms of action, biological pathways involved, and therapeutic effects, which we hypothesize can enhance the predictive power of DTI models.

The DTI prediction task is thus defined as learning a function or model ℱ that maps a drug–target pair, potentially enriched by their corresponding textual descriptions, to an interaction probability:

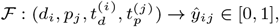

where *Ŷ*^*ij*^ represents the predicted probability of interaction between drug *d*^*i*^ and target *p*^*j*^.

### 3.2 The Proposed Method: DrugLM

We present a two-stage framework, **DrugLM**, which decouples the generation of LM embeddings from the actual DTI prediction task. This modular design provides two primary benefits: (1) it facilitates semantic alignment between drug and target representations via their textual descriptions, and (2) it introduces architectural flexibility, allowing seamless integration with diverse DTI prediction models. An overview of the framework is illustrated in Figure 1.

**Fig. 1:**
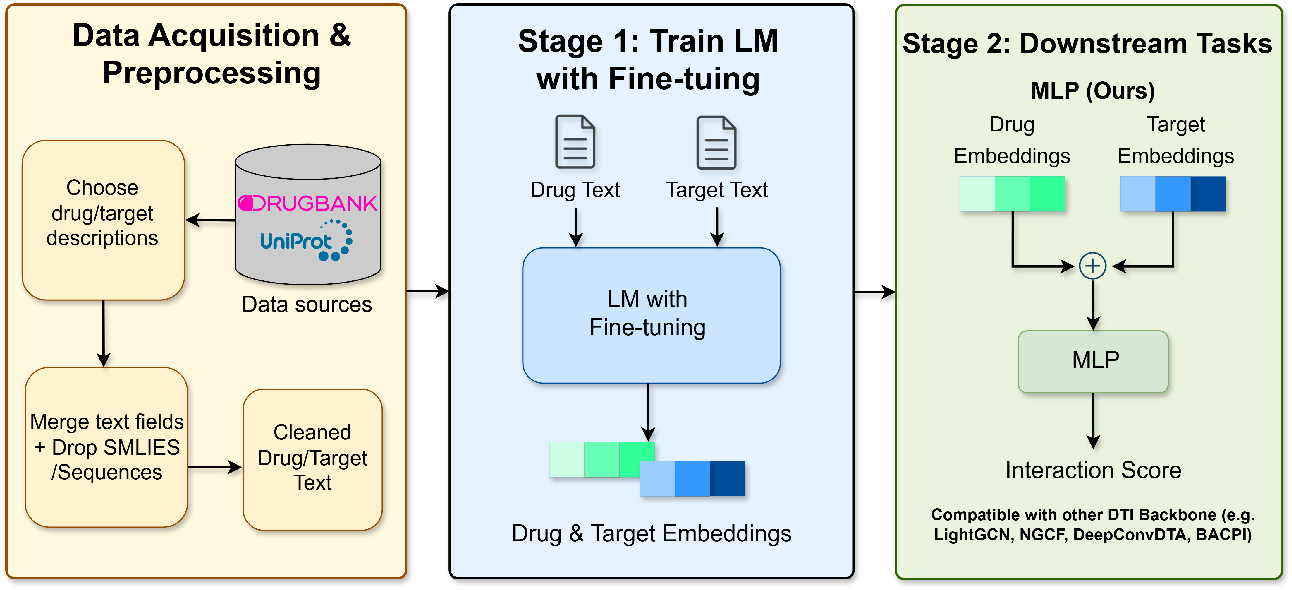
Overview of our two-stage framework for DTI prediction. We first collect drug-target interaction data along with textual descriptions from DrugBank. In Stage 1, we train a language model (with or without fine-tuning) to generate embeddings for drugs and targets based on their textual descriptions. In Stage 2, these textual embeddings are integrated into downstream models such as DeepConv-DTA, BACPI, GraphDTA, NGCF, LightGCN, or a simple MLP.

#### 3.2.1 Data Collection

We utilize DrugBank^2^, a comprehensive biomedical database containing detailed molecular, pharmacological, and interaction information for both approved and experimental drugs. In addition to molecular profiles, DrugBank provides rich annotations on drug–target interactions, therapeutic indications, and related biological pathways—making it a valuable resource for tasks such as DTI prediction, drug property prediction and bioactivity modeling.

For each drug, we extract its associated textual description and metadata, including therapeutic use, pharmacological mechanism, pharmacokinetic properties, and physicochemical characteristics, from DrugBank. To avoid information leaks, direct drug-target relationships were not included in the text description. For each target protein, we retrieve the corresponding natural language information from UniProt^3^. To ensure compatibility with general-purpose LMs, we apply the following preprocessing steps:

1. **Text Concatenation:** For each drug and each target, we concatenate all available textual description into a single unified text representation suitable for language model encoding.
2. **Exclusion of Non-Natural Language Elements:** Structured data such as SMILES strings (for drugs), InChI identifiers, amino acid sequences, and gene sequences (for proteins) are removed. These formats are not directly interpretable by general-purpose LMs and may degrade model performance if not appropriately handled.

As a result, each drug and each target is associated with a unified natural language description, collectively denoted as 𝒯^*d*^ for drugs and 𝒯^*p*^ for targets as defined earlier. These textual representations form the input for the subsequent LM-based embedding generation.

#### 3.2.2 Language Model Embeddings

Textual descriptions of drugs and targets encode rich biological information relevant to drug–target interactions. To harness this information, we employ two strategies for obtaining language model-based embeddings: (1) utilizing pre-trained models without fine-tuning, and (2) performing task-specific fine-tuning to better align representations with the DTI prediction objective.

##### Pre-trained LM Embeddings Without Fine-tuning

We begin by leveraging language models that have been pretrained on large-scale biomedical corpora. These models are assumed to encode deep semantic knowledge, including biomedical terminology, functional relationships and semantics. By directly applying these models, we extract embeddings from the natural language descriptions of drugs and proteins without requiring any additional parameter updates. Formally, given the textual descriptions 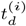 and 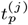 for drug *i* and protein *j*, and a pre-trained language model LM(*·*), the corresponding semantic embeddings are computed as:

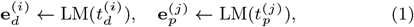

where 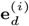 and 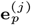 denote the embedding vectors for drug *i* and target *j*, respectively.

##### LM Embeddings with Fine-tuning

Fine-tuning a language model refers to the process of adapting a pre-trained model to a specific downstream task by continuing its training on a smaller, domain-specific dataset (Hu *et al*., 2022). In this work, we explore whether fine-tuning can enhance the ability of LMs to capture nuanced relationships between drugs and targets. We fine-tune the LM using a contrastive learning objective tailored to the DTI prediction task. Specifically, we minimize the contrastive cross-entropy loss:

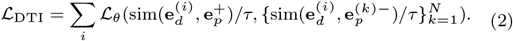

where sim(*·, ·*) denotes the cosine similarity between embeddings, 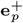 is the positive target embedding that interacts with drug *i*, and 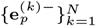 are negative (non-interacting) target embeddings.

Here, *τ* is the temperature parameter and *θ* represents the learnable parameters of the LM. We set *N* = 4: since the dataset contains only positive interaction pairs, we randomly sample four negative target embeddings for each positive pair from all non-interacting targets. We sum over *i* to aggregate the contrastive loss across all positive drug–target pairs. Although we treat all unknown pairs as negatives, one could further refine this by hard-negative mining or by excluding known off-target interactions. If a drug has multiple true targets, each (*d, p*) pair is treated separately in the above summation. After fine-tuning, the resulting embeddings are tailored to the DTI prediction task.

##### Fine-tuning Strategy

To ensure both computational efficiency and training stability, we adopt a supervised, parameter-efficient fine-tuning strategy. Specifically, we freeze the lower layers of the transformer, which primarily capture general linguistic patterns, and fine-tune only the upper layers that are more task-specific^4^. Formally, let the parameters of the language model be denoted by *θ*. We partition them as follows:

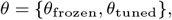

where *θ*^frozen^ corresponds to the parameters of the lower (frozen) layers, and *θ*^tuned^ denotes the parameters of the upper layers, which are fine-tuned for the DTI prediction task.

#### 3.2.3 Selection of Pretrained LMs

In this study, we select candidate language models from the MTEB leaderboard, prioritizing models that have been pretrained for semantic retrieval tasks. Considering factors such as model size, architecture, and performance on both retrieval and classification benchmarks, we primarily focus on three high-performing models: **BGE-large-en-v1.5, GTE-large-en-v1.5**, and **E5-large-v2**. BGE-large-v1.5 leverages RetromAE pre-training and large-scale contrastive fine-tuning on text pairs to alleviate similarity distribution imbalance and provide robust retrieval embeddings (Xiao *et al*., 2024). GTE-large-en-v1.5 builds on a Transformer++ encoder (BERT + RoPE + GLU) to support long-context embedding of up to 8 192 tokens, producing robust representations for extended input sequences (Zhang *et al*., 2024; Li *et al*., 2023). E5-large-en-v2 uses a 24-layer Transformer encoder pre-trained with weakly supervised contrastive learning on large-scale text pairs, yielding robust single-vector embeddings for both retrieval and classification tasks (Wang *et al*., 2022). Throughout the paper, we simply use BGE, GTE and E5 to denote these models.

##### Embedding Extraction Methods

Different LM architectures require distinct strategies for extracting embeddings. For models such as GTE and BGE, we extract the representation from the special classification token (<CLS>), which is a dedicated placeholder inserted at the very start of each input sequence, whose final-layer hidden state is trained to capture a holistic summary of the entire input. In contrast, for the E5 model, we apply average pooling over all token embeddings to obtain the final representation. Additionally, we apply the model-specific prompting templates recommended by each model’s developers to improve embedding quality. Specifically, for E5 we prepend “query: “to drug texts and “passage:” to target-protein descriptions; for BGE we add “Represent this sentence:” before every input (drugs and proteins alike); and for GTE we use the raw text directly, without any prefix.

#### 3.2.4 DTI Prediction Model Selection

Our proposed DrugLM framework is model-agnostic and can be seamlessly integrated into any downstream DTI prediction architecture. By leveraging language models, the resulting embeddings are expected to effectively capture semantic features aligned with biological mechanisms and functional properties relevant to drug–target interactions. These LM-derived embeddings are incorporated as complementary features to enhance the performance of various DTI prediction models. In this work, we primarily focus on five widely adopted model architectures for the DTI task. In addition, we also adopt a simple MLP model as another baseline to evaluate the performance improvement when incorporating the LM-derived embeddings. We briefly summarize the specific strategies we adopt for each of the DTI prediction models here.

- **DeepConv-DTI** (Lee *et al*., 2019): DeepConv-DTI takes the binary fingerprint representation 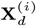 for drug *i* and the protein sequence 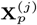 for target *j* as its inputs. It employs fully connected layers to learn a latent drug representation: 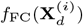 and adopts a convolutional neural network to detect localized residue patterns of target: 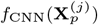. To incorporate textual information, we simply concatenate their original latent representations with LM-derived embeddings (from Eq. (1) or Eq. (2)) as follows:

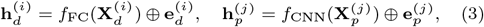

where 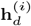 and 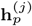 are the final representations of drug *i* and target *j*, respectively. These representations are concatenated and passed through fully connected layers to predict the interaction likelihood.
- **GraphDTA** (Nguyen *et al*., 2020): Similar to DeepConv-DTI, GraphDTA employs CNNs to learn protein sequence representations. However, instead of using fully connected layers for drug binary fingerprint, it models drugs as graphs and applies graph neural networks (GNNs) to learn their representations: 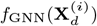. We integrate LM embeddings by concatenating them as follows:

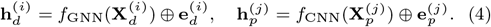
- **NGCF** (Wang *et al*., 2019) and **LightGCN** (He *et al*., 2020): Both approaches were initially developed for recommender systems to impute missing links between users and items. They can equally be applied to the DTI prediction problem by representing drug–target interactions as a bipartite graph and utilizing GNNs to learn embeddings for both drugs and targets by propagating information through the graph structure. Because their original work didnot consider the DTI application, we modify their implementations slightly by considering two alternatives for the node embedding initialization: (1) random initialization as a baseline, and initialization with the LM-based textual embeddings. This comparison allows us to assess the impact of semantic information on DTI prediction.
- **BACPI** (Li *et al*., 2022): BACPI employs a graph attention network for drugs and a convolutional neural network for targets. Following the same strategy as in DeepConv-DTI and GraphDTA, we augment the original representations with our text-based embeddings. These combined embeddings are then fed into a bidirectional attention network to model drug–target interactions.
- **MLP**: As a simple yet informative baseline, we propose a multilayer perceptron (MLP) model that directly utilizes our LM-based textual embeddings. Given the drug embedding 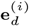 and target embedding 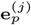, we concatenate them and use the MLP to predict the interaction:

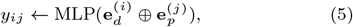

where *y*^*ij*^ denotes the predicted probability of interaction between drug *i* and target *j*. This setup allows us to evaluate whether semantic information alone is sufficient for effective DTI prediction.

## 4 Experiments

In this section, we evaluate the performance of DrugLM on real-world datasets. We curated a dataset consisting of 13,924 validated drug–target interactions, involving 6,673 unique drugs and 3,890 distinct targets. The data collection and preprocessing procedures are detailed in Section 3.2.1. To facilitate model training and evaluation, we employed a stratified edge-based splitting strategy: for drugs with fewer than 10 associated targets, all their edges were placed in the training set; for drugs with 10 or more targets, 80% of their edges were used for training and 20% for testing. Finally, 20% of the combined training edges were randomly held out as a validation set. The training set was used to train both the proposed models and the LM-based embeddings. The validation set was used for hyperparameter tuning and performance monitoring during training, while the test set was held out to ensure an unbiased evaluation of the final model performance.

To ensure fair comparison, SMILES-based baselines (DeepConv-DTI, GraphDTA, BACPI) were evaluated only on drugs with valid SMILES (excluding biologics), whereas LightGCN, NGCF and MLP were applied to both small molecules and biologics.

Following established protocols from prior works (Huang et al., 2020; Lee et al., 2019), we adopt some standard classification-based metrics for the binary prediction task. Specifically, we report three commonly used metrics: Accuracy (ACC), Area Under the Receiver Operating Characteristic Curve (AUC), and Area Under the Precision–Recall Curve (AUPR). To ensure robustness and reproducibility, all experiments were repeated five times with different random seeds. The final results are reported as the average of these five independent runs, providing a reliable estimate of the model’s stability and generalizability. Additionally, we performed paired two-sided t-tests on the five independent runs to assess statistical significance of embedding improvements.

### 4.1 Effectiveness of DrugLM

We present a comprehensive comparative analysis of various embedding strategies and model architectures for DTI prediction. Table 1 summarizes the overall results in terms of AUC, ACC, and AUPR. Our findings underscore the substantial impact by including language embeddings. Key observations are outlined below:

**Table 1.**
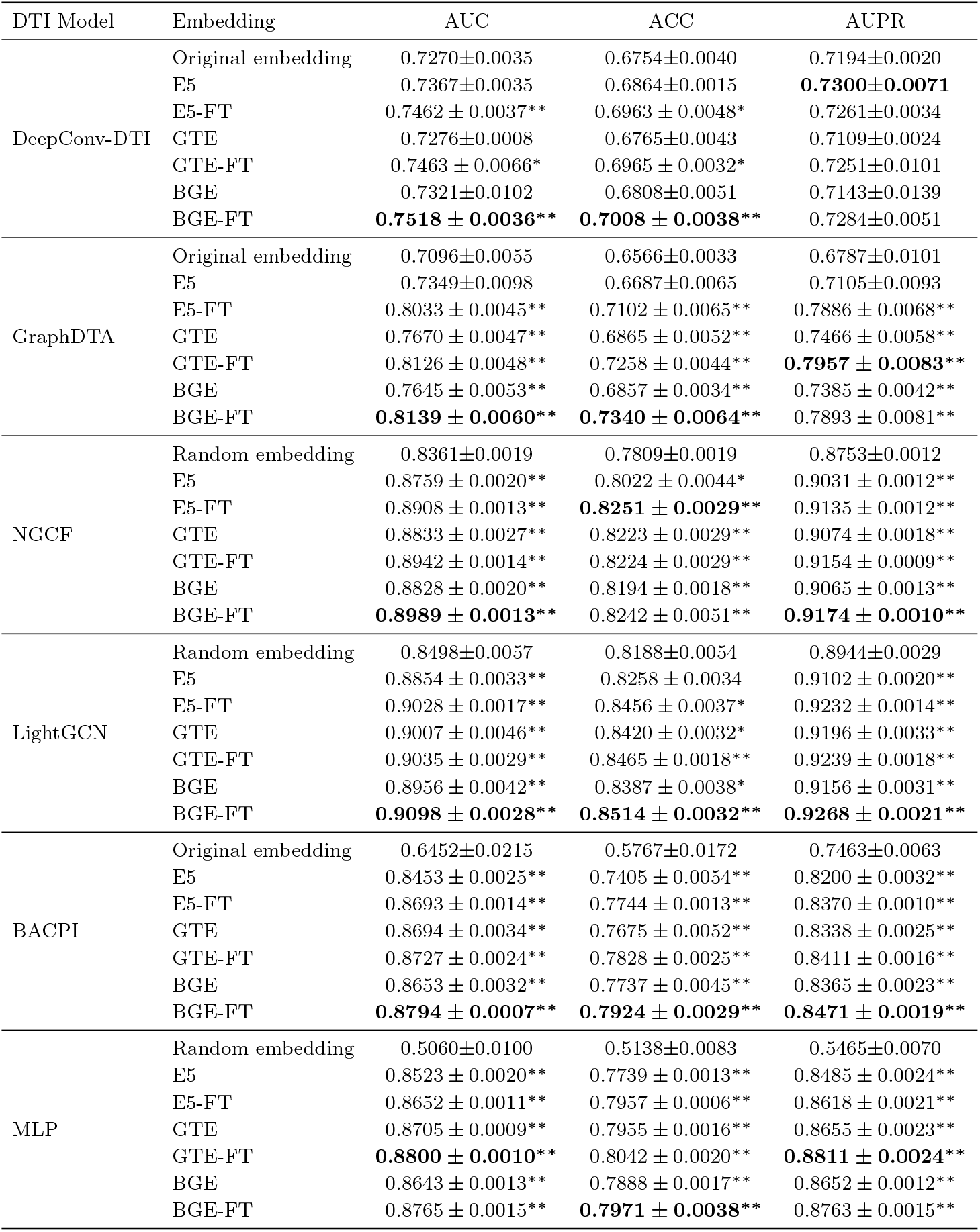
Performance of different language model embeddings on DTI prediction (mean±standard error over five independent runs). Bold values highlight the best results for each DTI model. FT: fine-tuned. Significance (paired t-test vs baseline): * *p <* 0.05, ** *p <* 0.01 (baseline = Original Embedding for DeepConv-DTI, GraphDTA, BACPI; Random Embedding for NGCF, LightGCN, MLP).

| DTI Model | Embedding | AUC | ACC | AUPR |
| --- | --- | --- | --- | --- |
| DeepConv-DTI | Original embedding | 0.7270±0.0035 | 0.6754±0.0040 | 0.7194±0.0020 |
|  | E5 | 0.7367±0.0035 | 0.6864±0.0015 | **0.7300±0.0071** |
|  | E5-FT | 0.7462 ± 0.0037** | 0.6963 ± 0.0048* | 0.7261±0.0034 |
|  | GTE | 0.7276±0.0008 | 0.6765±0.0043 | 0.7109±0.0024 |
|  | GTE-FT | 0.7463 ± 0.0066* | 0.6965 ± 0.0032* | 0.7251±0.0101 |
|  | BGE | 0.7321±0.0102 | 0.6808±0.0051 | 0.7143±0.0139 |
|  | BGE-FT | **0.7518 ± 0.0036\*\*** | **0.7008 ± 0.0038\*\*** | 0.7284±0.0051 |
| GraphDTA | Original embedding | 0.7096±0.0055 | 0.6566±0.0033 | 0.6787±0.0101 |
|  | E5 | 0.7349±0.0098 | 0.6687±0.0065 | 0.7105±0.0093 |
|  | E5-FT | 0.8033 ± 0.0045** | 0.7102 ± 0.0065** | 0.7886 ± 0.0068** |
|  | GTE | 0.7670 ± 0.0047** | 0.6865 ± 0.0052** | 0.7466 ± 0.0058** |
|  | GTE-FT | 0.8126 ± 0.0048** | 0.7258 ± 0.0044** | **0.7957 ± 0.0083\*\*** |
|  | BGE | 0.7645 ± 0.0053** | 0.6857 ± 0.0034** | 0.7385 ± 0.0042** |
|  | BGE-FT | **0.8139 ± 0.0060\*\*** | **0.7340 ± 0.0064\*\*** | 0.7893 ± 0.0081** |
| NGCF | Random embedding | 0.8361±0.0019 | 0.7809±0.0019 | 0.8753±0.0012 |
|  | E5 | 0.8759 ± 0.0020** | 0.8022 ± 0.0044* | 0.9031 ± 0.0012** |
|  | E5-FT | 0.8908 ± 0.0013** | **0.8251 ± 0.0029\*\*** | 0.9135 ± 0.0012** |
|  | GTE | 0.8833 ± 0.0027** | 0.8223 ± 0.0029** | 0.9074 ± 0.0018** |
|  | GTE-FT | 0.8942 ± 0.0014** | 0.8224 ± 0.0029** | 0.9154 ± 0.0009** |
|  | BGE | 0.8828 ± 0.0020** | 0.8194 ± 0.0018** | 0.9065 ± 0.0013** |
|  | BGE-FT | **0.8989 ± 0.0013\*\*** | 0.8242 ± 0.0051** | **0.9174 ± 0.0010\*\*** |
| LightGCN | Random embedding | 0.8498±0.0057 | 0.8188±0.0054 | 0.8944±0.0029 |
|  | E5 | 0.8854 ± 0.0033** | 0.8258 ± 0.0034 | 0.9102 ± 0.0020** |
|  | E5-FT | 0.9028 ± 0.0017** | 0.8456 ± 0.0037* | 0.9232 ± 0.0014** |
|  | GTE | 0.9007 ± 0.0046** | 0.8420 ± 0.0032* | 0.9196 ± 0.0033** |
|  | GTE-FT | 0.9035 ± 0.0029** | 0.8465 ± 0.0018** | 0.9239 ± 0.0018** |
|  | BGE | 0.8956 ± 0.0042** | 0.8387 ± 0.0038* | 0.9156 ± 0.0031** |
|  | BGE-FT | **0.9098 ± 0.0028\*\*** | **0.8514 ± 0.0032\*\*** | **0.9268 ± 0.0021\*\*** |
| BACPI | Original embedding | 0.6452±0.0215 | 0.5767±0.0172 | 0.7463±0.0063 |
|  | E5 | 0.8453 ± 0.0025** | 0.7405 ± 0.0054** | 0.8200 ± 0.0032** |
|  | E5-FT | 0.8693 ± 0.0014** | 0.7744 ± 0.0013** | 0.8370 ± 0.0010** |
|  | GTE | 0.8694 ± 0.0034** | 0.7675 ± 0.0052** | 0.8338 ± 0.0025** |
|  | GTE-FT | 0.8727 ± 0.0024** | 0.7828 ± 0.0025** | 0.8411 ± 0.0016** |
|  | BGE | 0.8653 ± 0.0032** | 0.7737 ± 0.0045** | 0.8365 ± 0.0023** |
|  | BGE-FT | **0.8794 ± 0.0007\*\*** | **0.7924 ± 0.0029\*\*** | **0.8471 ± 0.0019\*\*** |
| MLP | Random embedding | 0.5060±0.0100 | 0.5138±0.0083 | 0.5465±0.0070 |
|  | E5 | 0.8523 ± 0.0020** | 0.7739 ± 0.0013** | 0.8485 ± 0.0024** |
|  | E5-FT | 0.8652 ± 0.0011** | 0.7957 ± 0.0006** | 0.8618 ± 0.0021** |
|  | GTE | 0.8705 ± 0.0009** | 0.7955 ± 0.0016** | 0.8655 ± 0.0023** |
|  | GTE-FT | **0.8800 ± 0.0010\*\*** | 0.8042 ± 0.0020** | **0.8811 ± 0.0024\*\*** |
|  | BGE | 0.8643 ± 0.0013** | 0.7888 ± 0.0017** | 0.8652 ± 0.0012** |
|  | BGE-FT | 0.8765 ± 0.0015** | **0.7971 ± 0.0038\*\*** | 0.8763 ± 0.0015** |

- **Adding LLM embeddings significantly improve the performance of all models** Across all tested DTI prediction models, incorporating LLM embeddings (E5, GTE, BGE) consistently results in significant performance gains compared to the models using their original representations alone, or randomly initialized embeddings. The scales of improvements vary depending on the DTI models, the LMs used in generating the embedding, and pre-trained or fine-tuned embeddings. Across all the experiments and all the evaluation metrics, BGE with fine-tuning achieved the best results.
- **LLM embeddings with fine-tuning consistently boosts performance:** Fine-tuned LLM embeddings outperform their non-fine-tuned counterparts across various DTI models, reinforcing the effectiveness of domain adaptation via fine-tuning. For instance, using E5-FT with GraphDTA increases the AUC from 0.7349 to 0.8033 (a relative 9.31% gain) and the AUPR from 0.7105 to 0.7886 (a 10.99% gain) over non-fine-tuned E5.
- **GNNs benefit from bipartite drug–target graph topology:** Models that explicitly exploit the bipartite structure of drug–target interactions, such as NGCF and LightGCN, tend to outperform sequence-based and structure-based methods like DeepConv-DTI and GraphDTA, particularly when paired with high-quality embeddings. For example, LightGCN combined with BGE-FT achieves an AUC of 0.9098, substantially higher than DeepConv-DTI (0.7518) and GraphDTA (0.8139), highlighting the benefit of graph-based structural modeling in DTI tasks.
- **Potential knowledge conflict with LLM embeddings:** Interestingly, although fine-tuned embeddings generally enhance performance, some of them show a slight decline when used with certain DTI models such as DeepConv-DTI. For instance, E5-FT yields an AUPR of 0.7261, a 0.53% drop from the original E5’s AUPR of 0.7300. This suggests a potential conflict or redundancy between the contextual knowledge captured by LLMs and the handcrafted features (e.g., SMILES or amino acid sequences) utilized by the DeepConv-DTI architecture. Nonetheless, the general benefits of LLM embeddings remain substantial.
- **DrugLM significantly enhances even simple MLPs:** A particularly notable result is the strong performance of a simple multi-layer perceptron (MLP) when augmented with our LLM embeddings. Despite its simplicity, the MLP model achieves competitive and often superior results compared to more complex architectures. For example, MLP with GTE-FT achieves an AUC of 0.8811, outperforming DeepConv-DTI with BGE-FT (0.7463), GraphDTA with E5-FT (0.8033), and even BACPI with with GTE-FT (0.8727). This result demonstrates that high-quality language model embeddings can dramatically boost the performance of even basic models, offering a compelling and computationally efficient alternative for the DTI prediction task.

In summary, our experiments show that DrugLM, through the use of advanced language model embeddings for both drugs and targets, provides a powerful framework for improving the DTI prediction performance. The substantial gains observed across diverse prediction architectures, including the very simple MLP model, underscore the value of leveraging rich, context-aware textual representations.

### 4.2 Visualization of Drug Embeddings

To further investigate the effectiveness of LM embeddings, we compare the embeddings generated by LightGCN under three initialization strategies: **Random** initialization, **BGE-NonFT** (BGE pre-trained without fine-tuning), and **BGE-FT** (BGE fine-tuned). Figure 2 shows the t-SNE projections of these embeddings of the three cases. These plots show the node embeddings after LightGCN training, where initial vectors are refined via neighborhood aggregation on the drug–target graph. To generate them, we first obtained initial embeddings under each initialization, then trained LightGCN and applied t-SNE to the resulting node vectors (panels a–c). All maps include 6,238 small-molecule drugs (blue dots) and 435 biotech drugs (orange triangles). The labels whether a drug was a small molecule drug or a biotech drug were not included in the original text description.

**Fig. 2:**
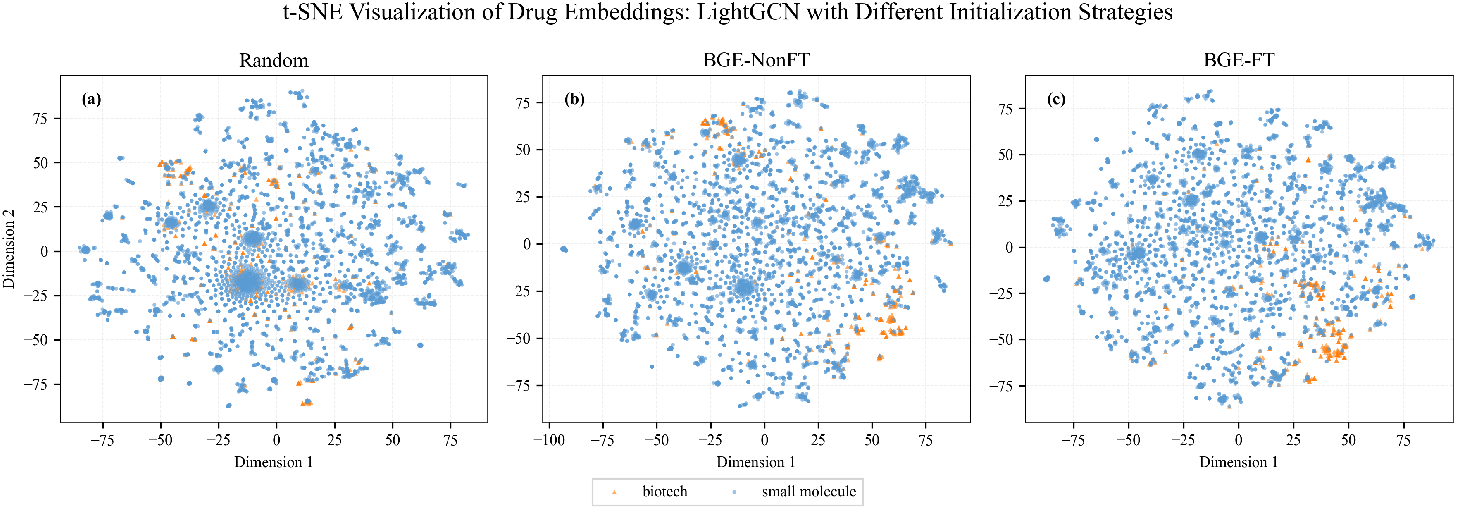
t-SNE visualization of drug embeddings generated by LightGCN under three initialization strategies: (a) **Random**, (b) **BGE-NonFT** (pre-trained BGE without fine-tuning), and (c) **BGE-FT** (fine-tuned BGE). Orange triangles indicate biotech drugs and blue circles indicate small-molecule drugs.

A key observation is that the final embeddings under the BGE initializations (BGE-NonFT and BGE-FT) exhibit much clearer separation between small-molecule and biotech drugs than those with random initialization. In panel (a), Random embeddings display substantial overlap between the two classes, whereas panels (b) and (c) show progressively distinct clusters. This pronounced clustering demonstrates that language models capture domain-specific information in the embedding space and that graph-based refinement further accentuates these semantic distinctions. Such well-separated groupings are critical for precise DTI prediction and underscore the value of LM-based representations combined with graph learning for accelerating drug discovery.

### 4.3 Training Effectiveness

To evaluate the training dynamics and effectiveness of DrugLM, we tracked the validation accuracy of all downstream DTI prediction backbones for 30 epochs under three initialization schemes: **Random/Original** initialization, **BGE-NonFT** (BGE pre-trained without fine-tuning), and **BGE-FT** (BGE fine-tuned). Figure 3 presents the validation accuracy curves for the LightGCN (a) and MLP (b) backbones. Analogous curves for DeepConv-DTI, GraphDTA, NGCF, and BACPI are provided in Supplementary Figure S1. These curves illustrate the downstream DTI training process rather than the language model pre-training phase. Note that for LightGCN and NGCF, validation metrics were recorded only every five epochs in their original implementations, whereas all other models report results at every epoch.

**Fig. 3:**
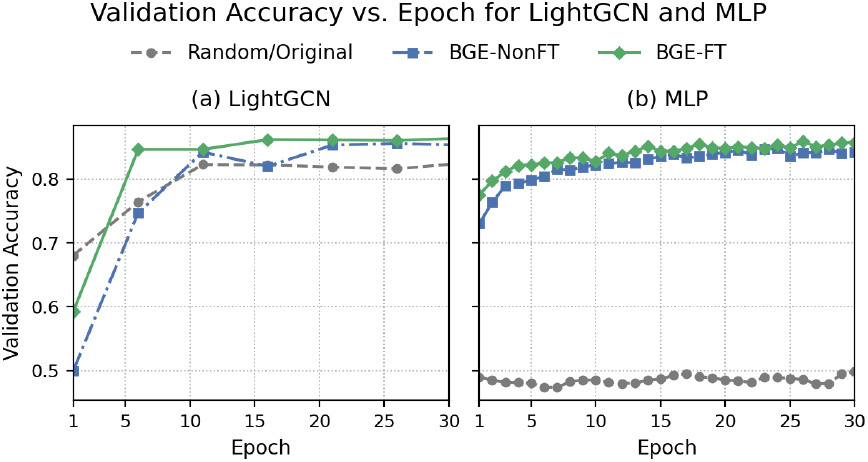
Validation accuracy across 30 epochs for LightGCN (a) and MLP (b). Three embeddings were used: **Random** initialization, **BGE-NonFT** (BGE pre-trained without fine-tuning), and **BGE-FT** (fine-tuned BGE).

Initializing DrugLM with pre-trained embeddings consistently yielded higher validation accuracy than random initialization for all models, highlighting the value of leveraging pretrained linguistic knowledge to create informative initial representations for drugs and targets. Fine-tuning the language model embeddings further amplified these gains by injecting additional domain-specific relevance, resulting in improved overall performance and more stable learning trajectories.

We also observed that the scale of improvements varied greatly for different model structures, which could be explained based on how the models actually worked. For example, when using random initialization for MLP, no meaningful information was provided as inputs. Therefore, the accuracy for the prediction was always around 0.5 for the binary classification. For LightGCN, the random initialization was only used for the node annotations of the drug-target bipartite graph. Although the annotations did not provide any information, the topological structure of the drug-target bipartite graph can still allow the model to predict meaningful results.

Notably, the magnitude of improvement varied substantially across different model structures, reflecting differences in how each model utilizes their own input representations. For instance, in the case of MLP, random initialization provided no meaningful input features, causing prediction accuracy to hover around 0.5 in this binary classification task. In contrast, for LightGCN, random initialization affected only the node features within the drug–target bipartite graph. Despite the lack of informative node annotations, LightGCN was still able to make meaningful predictions by leveraging the structural topology of the graph, demonstrating its relative robustness to feature initialization.

## Conclusion

We introduced **DrugLM**, a unified framework that advances drug-target interaction prediction by integrating rich, text-driven semantic information from language models into existing prediction model architectures. The core premise of DrugLM is that the extensive biomedical knowledge encoded in textual descriptions of drugs and targets—including mechanisms of action and therapeutic effects—can substantially enhance DTI prediction accuracy.

Comprehensive experiments demonstrated the effectiveness of DrugLM across various DTI prediction models. Leveraging LM-derived embeddings for drug and target representations consistently yielded significant performance gains. In particular, pre-trained LM embeddings outperformed random/original initialization, underscoring the value of linguistic prior knowledge. Moreover, parameter-efficient fine-tuning further improved both training efficiency and predictive performance by enhancing the domain-specific relevance of the embeddings. Remarkably, even a simple MLP model, when combined with LM embeddings, surpassed several specialized DTI architectures—highlighting the powerful semantic representation capabilities encoded in LMs.

Beyond the LM models discussed, we also evaluated embeddings generated by a larger autoregressive language model, *Llama3-7B*. While modest improvements were observed, they were limited—likely due to the causal attention mechanism inherent to autoregressive models, which constrains their ability to produce fully contextualized representations (Springer *et al*., 2025). We leave the investigation of even larger models, such as *Gemini* and *Llama3-80B*, to future work.

Overall, DrugLM offers a compelling and generalizable approach for incorporating advanced language models into DTI pipelines. By effectively leveraging unstructured biomedical text, DrugLM provides a simple yet powerful recipe for practical DTI prediction. The consistent benefits across diverse models and LMs suggest that DrugLM can substantially accelerate drug discovery by tapping into the wealth of semantic information embedded in biomedical literature.

## Supplementary Materials

### S1 Training Effectiveness Across Backbone Architectures and Different Embeddings

**Fig. S1:**
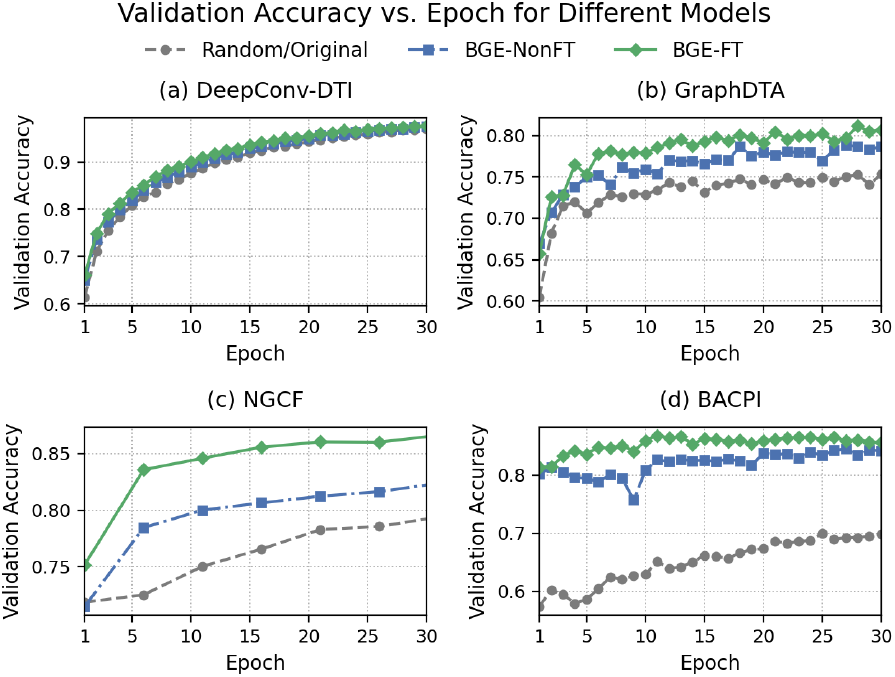
Validation accuracy across 30 epochs for different backbone architectures: (a) DeepConv-DTI, (b) GraphDTA, (c) NGCF, and (d) BACPI. For each model architecture, three embeddings were used: **Random/Original** initialization, **BGE-NonFT** (BGE pre-trained without fine-tuning), and **BGE-FT** (fine-tuned BGE).

## Footnotes

1 https://huggingface.co/spaces/mteb/leaderboard

2 https://go.drugbank.com/

3 https://www.uniprot.org/

